# Sensory population activity reveals confidence computations in the primate visual system

**DOI:** 10.1101/2024.08.01.606172

**Authors:** Zoe M. Boundy-Singer, Corey M. Ziemba, Robbe L. T. Goris

**Affiliations:** Center for Perceptual Systems, University of Texas at Austin, Austin, Texas, 78712, USA; McGovern Institute for Brain Research, Massachusetts Institute of Technology, Cambridge, MD, 02139, USA; Laboratory of Sensorimotor Research, National Eye Institute, National Institutes of Health, Bethesda, MD, 20892, USA

**Keywords:** neural coding, visual cortex, sensory uncertainty, population representation, metacognition

## Abstract

Perception is fallible^1–3^. Humans know this^4–6^, and so do some non-human animals like macaque monkeys^7–14^. When monkeys report more confidence in a perceptual decision, that decision is more likely to be correct. It is not known how neural circuits in the primate brain assess the quality of perceptual decisions. Here, we test two hypotheses. First, that decision confidence is related to the structure of population activity in sensory cortex. And second, that this relation differs from the one between sensory activity and decision content. We trained macaque monkeys to judge the orientation of ambiguous stimuli and additionally report their confidence in these judgments. We recorded population activity in the primary visual cortex and used decoders to expose the relationship between this activity and the choice-confidence reports. Our analysis validated both hypotheses and suggests that perceptual decisions arise from a neural computation downstream of visual cortex that estimates the most likely interpretation of a sensory response, while decision confidence instead reflects a computation that evaluates whether this sensory response will produce a reliable decision. Our work establishes a direct link between neural population activity in sensory cortex and the metacognitive ability to introspect about the quality of perceptual decisions.

## Introduction

Perceptual interpretations of the environment are automatically accompanied by a sense of confidence in this interpretation. For example, when soccer fans in a football stadium see a striker score a goal, they may hold their breath and ask other fans whether the ball really went in. Judging the trajectory of fast moving objects is difficult, and we know this. The ‘metacognitive’ ability to evaluate the quality of perceptual interpretations helps us to plan future actions^15^, learn from mistakes^16,17^, and optimize group decision-making^18^. How does the brain assess the quality of perceptual decisions? A prominent hypothesis is that early areas in sensory cortex provide raw sensory measurements which are used by downstream circuits in association cortex to guide perceptual decisions^19–22^ and assign confidence in these decisions^7,10,12,13^. It follows that there may exist a systematic relationship between neural population activity in sensory cortex and confidence in perceptual decisions. Here, we set out to test this hypothesis and document the basic characteristics of this relationship.

To examine the relationship between sensory activity and confidence, we developed a task that invites subjects to jointly report a perceptual decision and their confidence in this decision (also see the work by M. Vivar-Lazo & C.R. Fetsch, *SFN Abstract*, Population dynamics in areas MT and LIP during concurrent deliberation toward a choice and confidence report, 212.11, 2022). We trained two macaque monkeys (F and Z) to judge whether a visual stimulus presented near the central visual field was oriented clockwise or counterclockwise from vertical. The monkeys communicated their judgment with a saccade to one of four choice targets, organized in a rectangular pattern around the fixation mark (Fig. 1a). Horizontal saccade direction indicated the perceptual judgement, vertical saccade direction indicated the confidence in the decision. Choices were rewarded in a manner that incentivizes observers to introspect about decision quality on a trial-by-trial basis. Specifically, high confidence judgements resulted in a larger immediate reward when correct, but in a loss of potential future reward when incorrect (see Methods). While the animals performed this task, we recorded extracellular responses from neural populations in primary visual cortex (V1), the first sensory area in the primate visual system where individual neurons signal the task relevant feature, stimulus orientation^23^.

**Figure 1.**
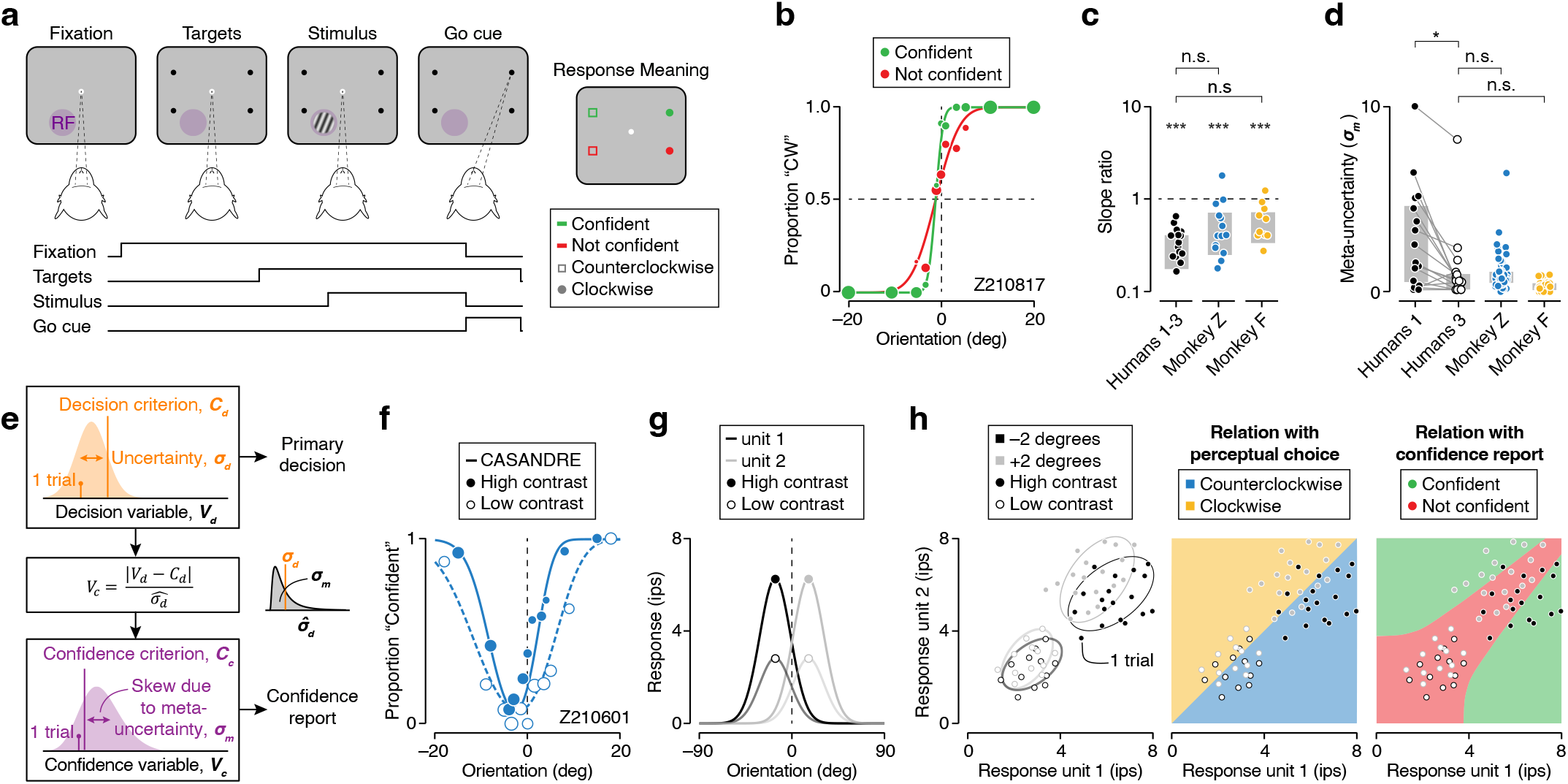
Perceptual confidence task: behavior and computational hypothesis. (**a**) Orientation discrimination task sequence. After the observer fixates for at least 500 ms, four choice targets appear, followed by the stimulus. The stimulus is placed in the neurons’ visual receptive field (RF). The observer judges whether the stimulus is rotated clockwise or counterclockwise relative to vertical. They jointly communicate this orientation judgment and their decision confidence with a saccade towards one of four choice targets. Horizontal saccade direction indicates the perceptual judgment, vertical saccade direction the confidence report. Correct decisions are followed by a juice reward (Methods). (**b**) Psychophysical performance for monkey Z in an example recording session. Proportion of clockwise (CW) choices for high-contrast stimuli is shown as a function of stimulus orientation, conditioned on the observer’s confidence report. Symbol size reflects the number of trials (total 527 trials, slope ratio = 0.57). The curves are fits of a behavioral model (Methods). (**c**) The ratio of the slope of high contrast psychometric functions. Sensible confidence judgments yield values smaller than one. Grey bars indicate interquartile range. (**d**) Meta-uncertainty for a group of human subjects and two monkeys. For the humans, each symbol represents metacognitive performance of one subject in one block of trials. For the monkeys, each symbol represent metacognitive performance in one behavioral session (humans: n = 17; monkey Z: n = 60; monkey F: n = 58). Grey bars indicate interquartile range. (**e**) Schematic of a process model for decision confidence^24^. (**f**) Proportion high confidence judgments as a function of stimulus orientation for high and low contrast stimuli (filled vs open symbols) in an example recording session. Symbol size reflects the number of trials (total 744 trials). Solid lines are fits of the process model shown in panel **e**. (**g**) Average firing rate as a function of stimulus orientation for two model neurons (black vs grey) and two stimulus contrasts (open vs closed symbols). (**h**) (Left) Joint responses of a pair of model neurons to repeated presentations of four stimuli that differ in orientation and contrast. (Middle) Illustration of a mapping rule that converts the pairwise activity into a perceptual decision. (Right) Illustration of a mapping rule that converts the same responses into a confidence report. Decision confidence is high when the sensory response is strong (towards upper right corner) and non-ambiguous (away from line of unity). n.s. not significant, * *P* < 0.05, ** *P* < 0.01, *** P < 0.001.

We found that confidence in perceptual decisions can be predicted from V1 population activity. The relationship between sensory activity and decision confidence appears as strong as the relationship between sensory activity and decision content. This assessment is based on the analysis of one hidden-layer neural networks trained to either predict the perceptual choice or the confidence report from V1 population activity. In both cases, the networks captured behavioral effects of stimulus manipulations (variations in stimulus orientation and stimulus contrast) as well as behavioral variability under repeated presentations of the same stimulus. As predicted by theoretical models of perceptual confidence, the relation between sensory activity and decision confidence fundamentally differs from the one between sensory activity and decision content. It involves an additional non-linearity and consideration of sensory uncertainty. Together, these results reveal how an essential metacognitive ability arises from downstream transformations of neural population activity in sensory cortex.

## Results

### Behavior and computational hypothesis

Both monkeys learned to report confidence in a fine orientation discrimination task. Their perceptual choices lawfully depended on stimulus orientation, and they made few errors in the easiest stimulus conditions (monkey F = ± 18.2 degrees, median performance, 100 % correct; monkey Z = ± 15.0 degrees, median performance, 100 % correct). Consider the choice behavior for an example recording session. Choices reported with high confidence are shown in green, choices reported with low confidence in red, and symbol size is proportional to the number of trials (Fig. 1b). As is evident from the raw data, for every stimulus condition, high confidence choices tended to be more accurate than low confidence choices (Fig. 1b, green vs red symbols). As a consequence, high confidence choices exhibited a steeper overall relationship with stimulus orientation (Fig. 1b, green vs red curve). We quantified this effect by estimating the slope of both psychometric functions and computing the slope ratio (Methods). For the vast majority of recording sessions, high confidence choices were associated with a steeper psychometric function than low confidence choices (monkey F: median slope ratio = 0.42, *P* < 0.001, Wilcoxon signed rank test against 1; monkey Z = 0.40, *P* < 0.001; Fig. 1c). This effect mirrored the choice behavior of a group of human subjects, naive to the purpose of our study, who performed a similar orientation discrimination task (*N* = 19, median slope ratio = 0.27, *P* < 0.001; Fig. 1c; Methods). These results suggest that the monkeys introspected about the quality of each perceptual decision and relied on a confidence assignment process that is qualitatively similar to the one used by humans (see Supp. Fig. 1 for further comparison).

We wondered whether the monkeys’ ability to assess the quality of perceptual decisions quantitatively resembles that of humans. The statistic we have considered thus far is inadequate to answer this question. The association between confidence and the slope of the psychometric function is a robust signature of sensible confidence assessments, but the slope ratio does not only depend on the quality of confidence assessments. It also depends on the subject’s perceptual sensitivity and their proclivity to report high confidence^24^. We therefore quantified the quality of the confidence reports by computing each subject’s meta-uncertainty (Methods). Meta-uncertainty is a statistic that expresses how well a decision maker can discriminate reliable from unreliable choices, regardless of their level of perceptual sensitivity and response biases^24^. Surprisingly, we found that the monkeys outperformed the human subjects during this group’s first visit to the lab (humans performed 1,100 trials in block 1, median *σ*_*m*_ = 1.56, median *σ*_*m*_ monkeys = 0.47, *P* < 0.001, Wilcoxon rank sum test; Fig. 1d, Humans 1 vs monkeys). We reasoned that task experience was the likely driver of this effect. To test this, we asked the human subjects to perform the experiment two more times. Reassuringly, they eventually caught up with the monkeys (median *σ*_*m*_ humans in block 3 = 0.57, median *σ*_*m*_ monkeys = 0.47, *P* = 0.58; Fig. 1d). We conclude that our animal paradigm invites high-quality metacognitive behavior.

What is the nature of the confidence assignment process that underlies this metacognitive capacity? Previous work has shown that choice-confidence data in tasks like ours are often well captured by a hierarchical process model in which confidence reflects an observer’s estimate of the reliability of their decision^24–27^. In these models, a stimulus gives rise to a noisy, one-dimensional decision variable (for example, a perceptual orientation estimate). Comparison of this decision variable with a fixed criterion yields a perceptual decision (‘clockwise’ or ‘counterclockwise’; Fig. 1e, top). The reliability of this decision is revealed by computing the distance between the decision variable and the decision criterion, and normalizing this distance by an estimate of the uncertainty of the decision variable (Fig. 1e, middle). Comparison of this decision reliability estimate with a fixed confidence criterion yields a confidence report (‘confident’ or ‘not confident’; Fig. 1e, bottom). The quality of the confidence reports is limited by a subject’s uncertainty about the uncertainty of the decision variable (‘meta-uncertainty’)^24^, or by an analogous noise term, depending on the specific model variant^25,26^. As can be seen for an example dataset, this computational framework captures how the monkey’s tendency to choose the ‘confident’ response option jointly depends on stimulus orientation and stimulus contrast (Fig. 1f, Supp. Fig. 1d,e).

Decision-making areas downstream of sensory cortex do not get one-dimensional perceptual estimates as input, but high-dimensional population responses. They implement operations akin to these idealized model computations by mapping this population activity onto the available choice options. To gain an intuition for these mapping rules, consider a pair of hypothetical V1 neurons whose responses selectively depend on stimulus orientation and stimulus contrast (Fig. 1g). One of these neurons prefers orientations smaller than 0 degrees, while the other one on average responds more vigorously to orientations larger than 0 degrees. Thus, their joint activity pattern contains information about stimulus orientation, regardless of stimulus contrast. Specifically, when neuron 2 is more active than neuron 1, the stimulus is more likely to be oriented clockwise from vertical and vice versa (Fig. 1h, left). The mapping rule used by a downstream decision-making circuit can thus be understood as projecting the population activity onto a one-dimensional axis perpendicular to a linear hyperplane that separates clockwise from counterclockwise response patterns (Fig. 1h, middle). The resulting decisions will not be flawless – due to neural response variability, there is considerable overlap between both response distributions, making errors inevitable. Crucially, the population response also contains information about the probability of such an error. The closer the population activity is to the hyperplane, the more probable an error. This effect is amplified for activity patterns that reside close to the bottom left corner of this state space. This part of the space is visited when the stimulus-drive is weak, for example because stimulus contrast is low or stimulus size is small. Here, response patterns are dominated by spontaneous activity, resulting in high levels of sensory uncertainty^28–31^ and many incorrect decisions (Fig. 1h, right). These geometrical considerations yield two testable predictions. First, that decision confidence is related to the structure of population activity in sensory cortex. And second, that this relation differs from the one between sensory activity and decision content.

### Predicting perceptual decision confidence from V1 activity

While the animals performed the perceptual confidence task, we used multilaminar electrode arrays to record population activity from ensembles of V1 units whose receptive fields overlapped with the stimulus location (Methods). Populations ranged in size from 8 to 46 units (median = 15 units). Consider the activity of three simultaneously recorded units. Stimulus onset elicited a strong transient response, followed by a weaker sustained response (Fig. 2a). Some units were better driven by counterclockwise orientations (Fig. 2a, top), some by clockwise orientations (Fig. 2a, bottom), and some did not differentiate between these stimulus conditions (Fig. 2a, middle). We first asked whether the observed populations could in principle provide the sensory signals to support the perceptual task. To this end, we trained linear *stimulus decoders* to discriminate between clockwise and counterclockwise stimuli and tested them on non-ambiguous hold-out trials (Methods; Fig. 2b). We found that each recorded population could support the perceptual task above chance level (neural performance ranged from 57.4 to 96.9% correct, median = 69.2%). These decoders have only been provided with neural population responses and stimulus labels (‘clockwise’ or ‘counterclockwise’). Yet it is natural to ask whether they can offer some insight into the monkeys’ behavior. We compared the stimulus decoders’ choices with the animals’ reports on a trial-by-trial basis and found that the fraction of correctly predicted perceptual decisions exceeded the number expected by chance (median difference = 6.3%, *P* < 0.001, Wilcoxon signed rank test; Fig. 2b). Both variables exhibited a clear relationship; the better the neural populations could support the perceptual task, the better the stimulus decoder predicted perceptual decisions (Pearson correlation: *r* = 0.79, *P* < 0.001, Fig. 2b). This finding is consistent with the hypothesis that decision-making circuits downstream of visual cortex estimate the most likely interpretation of a sensory response, just like these stimulus decoders do. We also compared the stimulus decoders’ output with the animals’ confidence reports and found them to be unrelated (median Pearson correlation = –0.03, *P* = 0.16; Fig. 2c). This result is not surprising. It simply confirms that in our task, the relationship between sensory activity and decision content cannot account for decision confidence.

**Figure 2.**
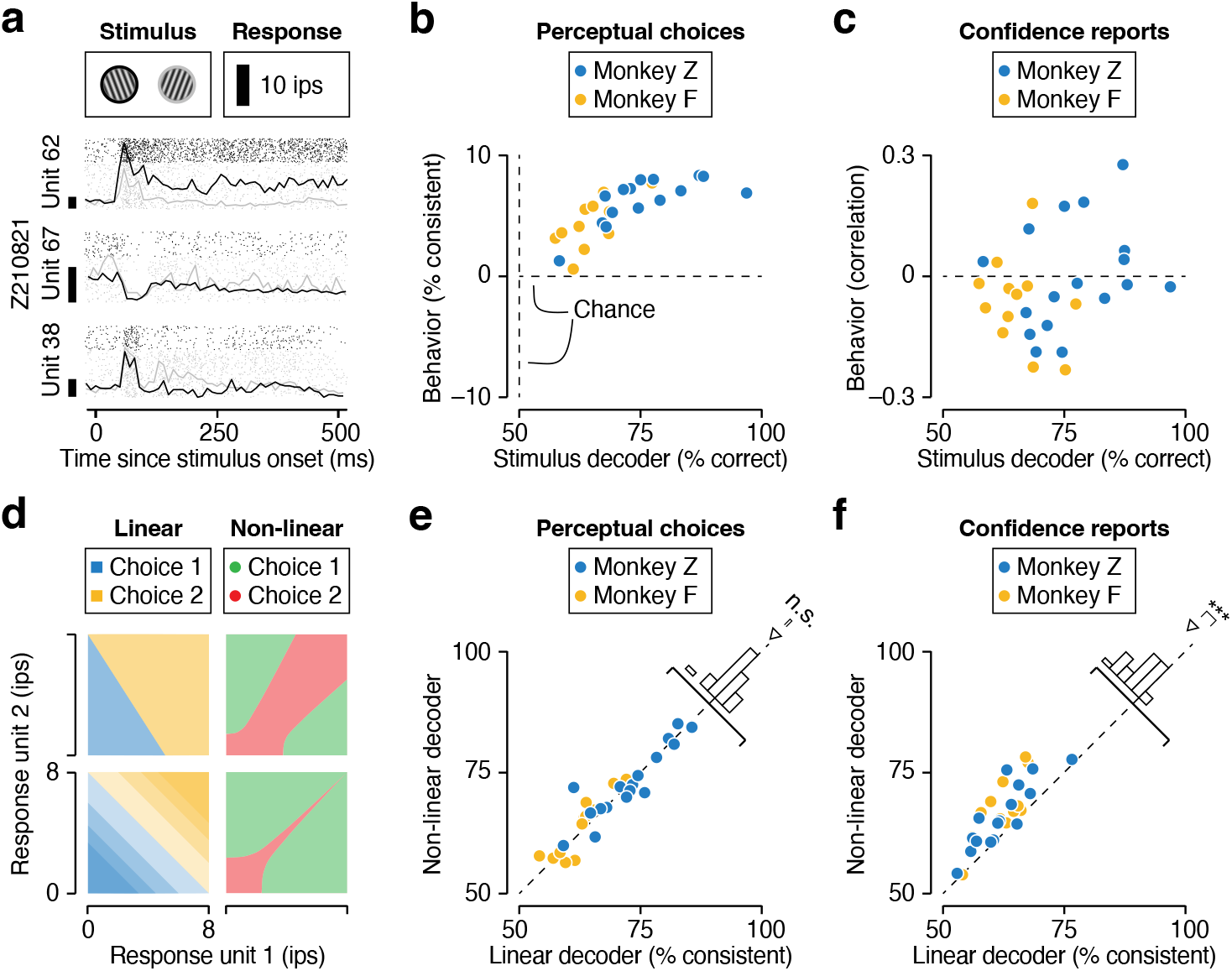
Predicting perceptual decisions and decision confidence from V1 population activity. (**a**) Spike rasters (dots) and PSTHs (lines) of three example units during presentation of a clockwise (gray) and counterclockwise (black) stimulus. (**b**) Analysis of the linear stimulus decoder. The proportion of correctly predicted perceptual choices minus the proportion expected by chance is plotted against the decoder’s task performance. Each symbol represents a recording session. (**c**) The correlation between the monkeys’ confidence reports and the linear stimulus decoder’s output is plotted against the decoder’s task performance. (**d**) Example mapping rules that can be implemented by a linear (left) and non-linear (right) decoder. (**e**) Comparison of the proportion correctly predicted perceptual choices by a linear (abscissa) and non-linear (ordinate) choice decoder. (**f**) Comparison of the proportion correctly predicted confidence reports by a linear (abscissa) and non-linear (ordinate) confidence decoder. n.s. not significant, * *P* < 0.05, ** *P* < 0.01, *** P < 0.001.

To expose the relationship between sensory population activity and decision confidence, we trained *confidence decoders* to discriminate between trials in which monkeys reported choices with either high or low confidence. For this analysis, we considered both linear and non-linear decoders (specifically, one-hidden layer neural networks; Methods). Linear decoders can slice a high-dimensional space in various ways (Fig. 2d, left), but none of the possible variations fully captures the hypothesized confidence mapping rule (Fig. 1h, right). Non-linear decoders can implement more complex input-output relations (Fig. 2d, right), making them better suited to test the hypothesis. Importantly, this additional complexity is not guaranteed to be beneficial. This will only be the case if the brain’s mapping rule requires the complexity (compare Fig. 1h, middle and right). To connect this concept to our data, we first compared linear and non-linear *choice decoders* trained to predict perceptual decisions (Methods). We orthogonalized the neural choice and confidence subspaces by curating the decoders’ training sets such that the animals’ perceptual choice contained no information about their confidence report and vice versa (Methods; Supp. Fig. 2a). As expected, linear and non-linear choice decoders performed similarly well, suggesting that a linear mapping rule suffices to relate sensory population activity to perceptual decisions (median performance linear choice decoder = 66.8% correctly predicted choices; non-linear choice decoder = 68.8%, median difference = -0.7%, *P* = 0.34, Wilcoxon signed rank test; Fig. 2e). We then performed the same comparative analysis on the confidence reports and obtained a different result. The nonlinear confidence decoders consistently outperformed their linear counterparts in predicting confidence (median linear confidence decoder = 61.8% correctly predicted confidence reports; non-linear confidence decoder = 65.6%, median difference = 4.0%, *P* < 0.001; Fig. 2f). In general, confidence reports could be predicted about as well as perceptual decisions (median performance difference between non-linear choice and confidence decoders = 3.5%, *P* = 0.19). Together, these results confirm that perceptual decision confidence is related to the structure of population activity in sensory cortex, and that this relationship is more complex than the relation between this activity and decision content.

### Interrogating the confidence decoder

We seek to understand how sensory population activity informs confidence in perceptual decisions. So far, our analysis suggests that non-linear decoders trained to predict behavioral choice-confidence reports from neural population activity are a powerful tool in this endeavour. Of course, this is only true to the extent that the mapping relation learned by the decoders resembles the one used by the brain. This need not be the case. Clearly, the confidence decoders are imperfect predictors of the animals’ behavior. It is possible that their success is based on exploiting idiosyncratic relationships between neural responses and confidence reports that are distinct from the brain’s confidence computation^32^. If this were the case, the confidence decoders’ output should not exhibit the key signature of sensible confidence assignments, nor should they be able to generalize to new testing conditions. We investigated both issues. We first computed the slope of the psychometric function, conditioned on the confidence decoder’s output (Methods; Fig. 3a). Higher confidence outputs were associated with a steeper psychometric function (median slope ratio = 0.88, *P* = 0.02, Wilcoxon signed rank test, Fig. 3b). This pattern recapitulates a key feature of the animals’ behavior and implies that the confidence decoder’s outputs are sensible. The confidence decoder recognizes which neural responses are more likely to result in a reliable perceptual decision.

**Figure 3.**
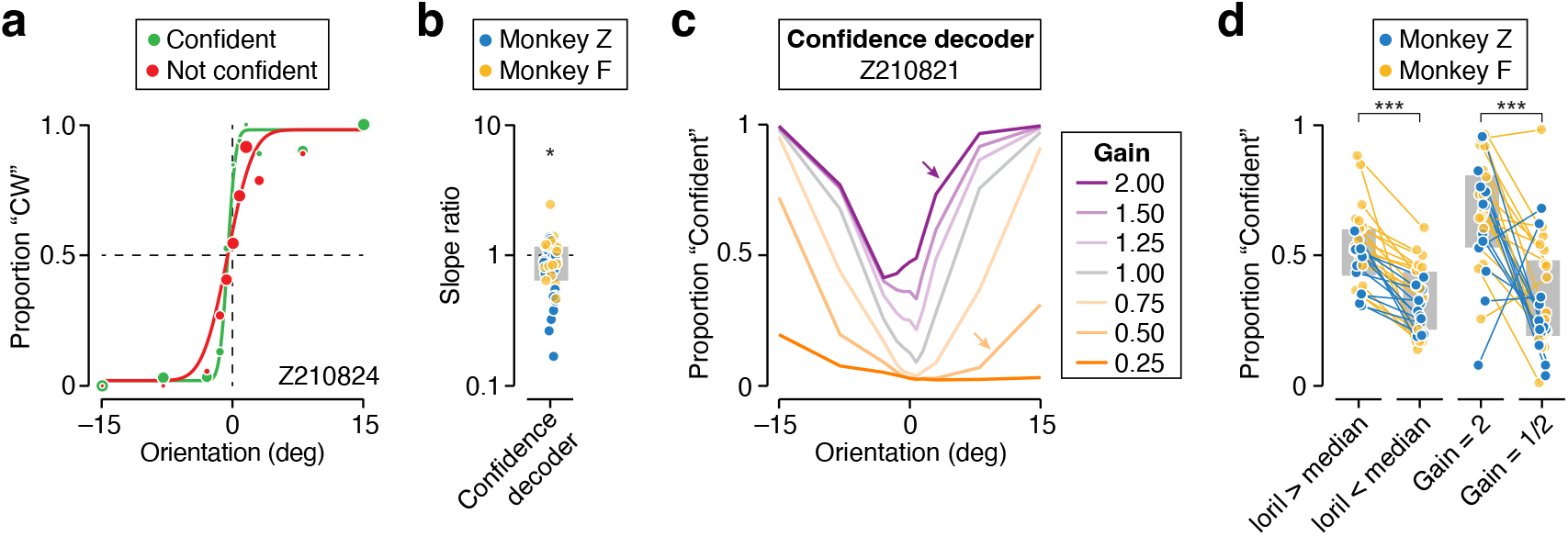
The non-linear confidence decoder yields sensible and robust outputs. (**a**) Psychophysical performance for monkey Z in an example recording session. Proportion of clockwise (CW) choices for high-contrast stimuli is shown as a function of stimulus orientation, conditioned on the confidence decoder’s output. Symbol size reflects the number of trials (total 520 trials, slope ratio = 0.88). The curves are fits of a behavioral model. (**b**) The ratio of the slope of both psychometric functions. Grey bars indicate interquartile range. (**c**) Illustration of the confidence decoder’s output for various synthetic patterns of neural activity for an example recording session. (**d**) Summary of the synthetic confidence experiments for all recording sessions. (Left) Proportion of predicted high confidence outputs elicited by stimuli whose orientation is more or less extreme than the median stimulus orientation. (Right) Proportion of high confidence outputs elicited by sensory input patterns with a high or low response gain (indicated by the colored arrows in panel **c**). Grey bars indicate interquartile range. n.s. not significant, * *P* < 0.05, ** *P* < 0.01, *** P < 0.001.

If the mapping rule learned by the confidence decoder resembles the one used by the brain, it should transfer to more challenging testing conditions, such as input patterns it has not been exposed to during training. To test this, we probed the confidence decoders with synthetic patterns of neural activity. We designed these patterns such that they would expose the ‘pure’ effects of stimulus orientation and stimulus contrast on decision confidence. Specifically, for every stimulus orientation, we created a synthetic pattern by computing the trial-averaged population response, thus removing the effects of neural response variability (Methods). We captured the effects of stimulus contrast by changing the gain of the synthetic neural responses^33–35^. Here, we went far outside the range of our experimental stimulus manipulation to create out-of-distribution inputs (Methods). Consider the decoder’s confidence-outputs for an example recording session. More extreme orientations are always associated with more high confidence outputs, regardless of the level of response gain (Fig. 3c). Additionally, higher levels of response gain are always associated with more high confidence outputs, regardless of the stimulus orientation (Fig. 3c). These effects were evident across datasets (median difference in predicted proportion high confidence outputs for more vs less extreme stimulus orientations = 15%, *P* < 0.001; median difference for a response gain of 0.5 and 2 = 38%, *P* < 0.001; Wilcoxon signed rank test; Fig. 3d). Thus, the decoder’s confidence output jointly depends on stimulus orientation and stimulus contrast, thereby recapitulating the second key feature of the animals’ behavior (Supp. Fig. 1e). Moreover, the mapping rule learned by the decoder generalizes to new testing conditions. We conclude that the confidence decoder evaluates neural activity in a sensible and robust manner.

### Relationship between choice and confidence computations

Our analysis of neural activity was inspired by a computational framework in which confidence reflects an observer’s estimate of the reliability of their decision^24–26^. In this framework, the computations that form a decision are distinct from the ones that assign confidence in these decisions. However, there is a direct relationship between the latent variables that underlie the overt perceptual choices and confidence reports. Specifically, more extreme decision variable values will yield higher confidence variable values (Fig. 1e, middle). The decoders we trained on neural data use a latent variable to predict behavioral choice-confidence reports (Methods). We wondered whether these latent variables would be related as predicted by the computational framework. If this were the case, it would provide direct evidence for the notion that the brain’s confidence computation evaluates the quality of the sensory evidence that informed the decision.

Consider the neurally decoded decision variable for an example recording session. There are three important effects. The decision variable varies linearly with stimulus orientation (Fig. 4a). The slope of this relationship depends on stimulus contrast (Fig. 4a, left vs right panel). And trials that culminate in a “clockwise decision” are associated with a higher decision variable value (Fig. 4a, yellow vs blue). These effects were present in most of our datasets (median slope for high contrast stimuli = 0.064, *P* < 0.001; median reduction in slope for low contrast stimuli = 0.02, *P* = 0.04; median change in offset with perceptual choice = 0.22, *P* < 0.001, Wilcoxon signed rank test; Fig. 4b). The neurally decoded confidence variable exhibits a different structure. It varies parabolically with stimulus orientation (Fig. 4c). The width and offset of the parabola depend on stimulus contrast (Fig. 4c). And trials that culminate in a “high confidence” report are associated with a higher confidence variable value (Fig. 4c). Again, these effects were present in most of our datasets (median quadratic coefficient for high contrast stimuli = 0.002, *P* < 0.001; median change in this coefficient for low contrast stimuli = 0.001, *P* = 0.005; median change in offset for low contrast stimuli = 0.15, *P* < 0.001; median change in offset with confidence report = 0.11, *P* < 0.001 Wilcoxon signed rank test; Fig. 4d). To compare the strength of the association of both decision and confidence latent variables with the overt behavior, we computed their ability to discriminate behavior in the absence of stimulus variation (Methods). A discriminability value of 50% corresponds to chance performance, while 100% means that the behavior can be perfectly predicted from the latent variable. For both perceptual choices and confidence reports, we found a modest association (median decision variable discriminability = 54%, *P* < 0.001; median confidence variable discriminability = 56%, *P* < 0.001, Wilcoxon signed rank test; Fig. 4b,d). This association tended to be stronger for populations that contained more stimulus information (Supp. Fig. 2b). Thus, the decoders’ latent variables provide insight into the neural processes underlying the overt choice-confidence reports.

**Figure 4.**
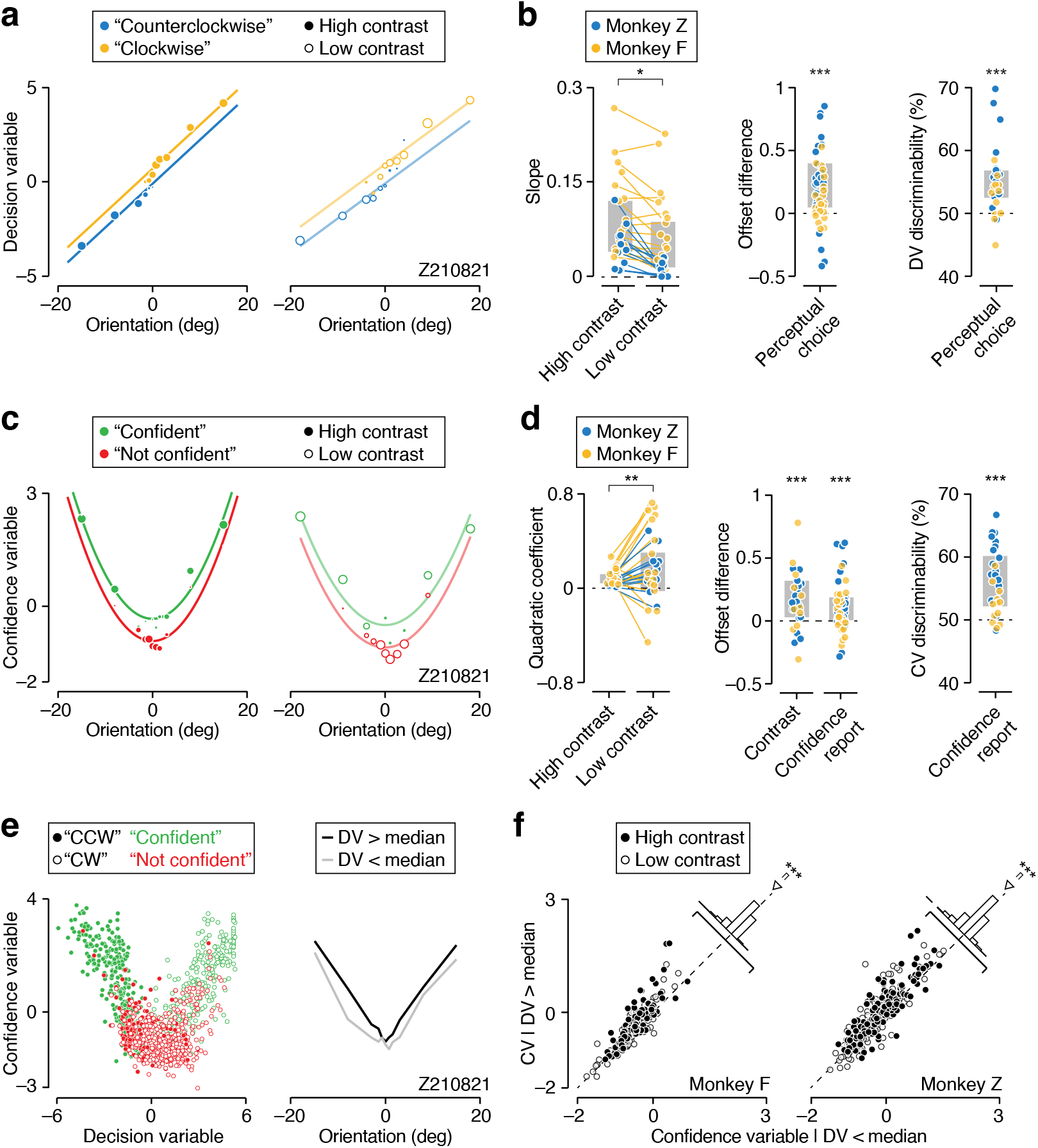
The decoders’ latent variables follow the predictions of a computational framework of perceptual decision confidence. (**a**) The latent variable of the nonlinear choice decoder plotted against stimulus orientation, for high and low contrast trials (left vs right), conditioned on the animal’s perceptual choice (yellow vs blue) for an example dataset. (**b**) Summary of the decision variable’s statistical structure across all recording sessions. Grey bars indicate interquartile range. (**c**) The latent variable of the nonlinear confidence decoder plotted against stimulus orientation, for high and low contrast trials (left vs right), conditioned on the animal’s confidence report (red vs green) for an example dataset. (**d**) Summary of the confidence variable’s statistical structure across all recording sessions. Grey bars indicate interquartile range. (**e**) Direct comparison of the confidence variable and the decision variable. (Left) Each point represents a single trial in an example recording session. (Right) The mean confidence level plotted against stimulus orientation for trials with a decision variable value that is more (black) or less (gray) extreme than the stimulus-specific median. (**f**) Summary of the median-split analysis illustrated in panel **e** for all recording sessions. Each data point represents a single stimulus condition. n.s. not significant, * *P* < 0.05, ** *P* < 0.01, *** P < 0.001.

Plotting the confidence variable against the decision variable for all trials of an example recording session reveals a U-shaped relationship, consistent with the proposed confidence computation (Fig. 4e, left). If this relationship arises from this computation, it should leave a signature even within the same stimulus conditions. Specifically, trials that yield a more extreme decision variable value should result in a higher value of the confidence variable. To test this prediction, we computed the median of the decision variable for every stimulus condition and computed the average of the confidence variable separately for trials above and below the median. As can be seen for an example recording session, more extreme decision variable values were systematically associated with higher levels of confidence (Fig. 4e, right). This effect was evident across all datasets, for both monkeys (median difference in confidence variable: monkey F = 0.06, *P* < 0.001, monkey Z = 0.11, *P* < 0.001; Fig. 4f). Our analysis ensured that choice and confidence signals occupied orthogonal neural dimensions (Methods). We therefore conclude that the brain’s confidence computation evaluates the same sensory population activity that informed the decision.

## Discussion

In this study, we investigated neural population activity in V1 during a perceptual confidence task. We sought to understand the brain’s confidence assignment process. This process underlies the metacognitive ability to evaluate the quality of perceptual interpretations. We suggest that confidence arises from a nonlinear transformation of the same sensory signals that inform perceptual decisions. When the sensory population response is strong and unambiguous, this transformation results in high decision confidence (Fig. 1h, green zone). Conversely, when the sensory population response is weak or ambiguous, it results in low levels of confidence (Fig. 1h, red zone). Our proposal is supported by three distinct observations. First, nonlinear decoders of V1 population activity can predict monkeys’ confidence in perceptual orientation judgments (Fig. 2f), establishing a direct link between the structure of sensory activity and decision confidence. Second, these decoders yield sensible and robust confidence outputs when presented with synthetic patterns of neural activity (Fig. 3), suggesting they capture the essence of the brain’s confidence computation. Third, trials that yield stronger and less ambiguous V1 responses as evidenced by a neurally decoded decision variable also result in higher levels of neurally decoded confidence (Fig. 4).

Our experimental paradigm enabled us to compare choice and confidence decoders trained on the same neural responses. We found that the relationship between sensory activity and decision confidence is as strong as the relationship between sensory activity and perceptual choice (Fig. 2e,f and Fig. 4b,d). However, we suggest that there is a fundamental difference between both relationships. Perceptual choices arise from a neural computation downstream of sensory cortex that identifies the most likely interpretation of the sensory response; decision confidence instead arises from a computation that evaluates whether this sensory response will produce a reliable decision (Fig. 1e). Because these computations are distinct, they can manifest as mapping rules of sensory population activity that occupy orthogonal neural subspaces (Fig. 1h). Previous work offers indirect support for these ideas. Specifically, micro-stimulating sensory neurons in a post-decision wagering task altered monkeys’ optout choice behavior as if they experienced a change in the sensory signal^10^. These results suggest that the same sensory signals that inform decision content inform decision confidence. A different study employing the postdecision wagering paradigm found that pulvinar neurons represent decision confidence but not perceptual choice^9^. Inactivating the pulvinar altered monkeys opt-out choice behavior but not their perceptual sensitivity^9^. These results suggest that distinct brain circuits may be responsible for decision formation and confidence assignment. Consistent with this, studies that employed a post-decision time investment task found that orbitofrontal cortex neurons in rats play a similar role^12,13^ and represent an abstract decision confidence signal^14^. Our work clarifies how neural circuits can extract such pure decision confidence signals from sensory population activity. However, note that in some experimental paradigms, the same neurons may represent decision content and confidence^7,10^.

We have shown that decoders of V1 activity capture behavioral effects of stimulus manipulations as well as behavioral variability under repeated presentations of the same stimulus (Fig. 4a,c). Correlations between between neural and behavioral responses can illuminate their causal relationship. However, these correlations can also arise for spurious reasons. Previous studies employing binary perceptual decision-making tasks found that choice-related signals in sensory cortex reflect a combination of factors. These include the perceptual decision-making process^36–39^, but also choice-aligned fluctuations in attention^40,41^, expectation^42^, motor planning^43^, and in other unspecified sources that impact sensory activity^44,45^. Could spurious reasons underlie the association between neural activity and overt behavior in our study? This concern is warranted. There is no statistical guarantee that the associations we reported primarily reflect the confidence assignment process. However, our task-design has a unique strength compared to binary decision-making tasks. Subjects generated a two-dimensional choice-confidence report. Our analysis ensured that both dimensions were orthogonal in neural population space. Nevertheless, we found that trials that yielded a more extreme perceptual decision variable also resulted in a higher level of confidence (Fig. 4e,f). For this association to arise for spurious reasons, there would need to be a factor whose properties are more complex than simple choice-alignment. It would need to jointly align with perceptual choices and confidence reports. While we cannot rule out this possibility, we hope that the richness of our behavioral paradigm has helped to expose the confidence computations implemented by neural circuits downstream of sensory cortex.

The ability to recognize which perceptual interpretations of the environment are at risk of being flawed is a hallmark of metacognition and as such often associated with higher intelligence. Our findings suggest that the confidence computations underlying this ability in the primate brain at least in part arise from simple deterministic transformations of sensory population activity. These transformations can in principle be realized in basic neural circuits, calling into question the extent to which confidence-mediated behavior truly provides insight into high level cognitive processes^46^. In general, metacognitive judgements are imperfect^24,25,47^. Here, this was evident from the levels of meta-uncertainty displayed by our human and non-human subjects (Fig. 1d). As of yet, we do not know the neural causes of this. Metacognitive inefficiencies may originate in noise in sensory representations^29,48–50^. Alternatively, these inefficiencies may arise downstream of sensory cortex, for example from sub-optimal confidence mapping rules^51^. The task-paradigm and computational framework we have developed offer promising vehicles to address these outstanding questions and achieve a more complete understanding of the neural mechanisms that underlie and constrain our sense of confidence.

## METHODS

### Animal subjects

Our experiments were performed on two adult male macaque monkeys (*Maccaca mulatta*, aged 7 and 10 years old at the time of the experiments). The animals were trained to perform an orientation discrimination task with saccadic eye movements as operant responses. Monkey F had previously participated in another research study^22^, Monkey Z had not previously participated in research studies. All training, surgery, and recording procedures were approved by the University of Texas Institutional Animal Care and Use Committee and conformed to the National Institutes of Health Guide for the Care and Use of Laboratory Animals. Under general anesthesia, both animals were implanted with three custom-designed titanium head posts and a titanium recording chamber which enabled access to V1^52^.

### Apparatus

The monkeys were seated in a custom-designed primate chair in front of a gamma-corrected 22-inch CRT monitor (Sony Trinitron, model GDM-FW900), with their heads restrained using three surgical implants. Stimuli were shown on the CRT monitor, which was positioned approximately 60 cm away from the monkeys’ heads. The CRT had a resolution of 1280 by 1024 pixels with a refresh rate of 75 Hz. Eye position was tracked continuously with an infrared eye tracking system at 1 kHz (EyeLink 1000, SR Research). Stimuli were presented using the Psychophysics Toolbox^53^ in MATLAB (MathWorks). Neural activity was recorded using the Plexon OmniPlex System (Plexon). Precise temporal registration of task events and neural activity was obtained through a Datapixx system (Vpixx). All of these systems were integrated using the PLDAPS software package^54^ (https://github.com/HukLab/PLDAPS). An analogous setup was used for the human psychophysical experiment, except that head position was stabilized using a chin rest and the monitor was a Hewlett Packard, model A7217.

### Visual stimuli

We constructed oriented visual stimuli by bandpass filtering 3-D luminance noise. The filter was orientation-spatial frequency-temporal frequency separable. All stimuli were constructed with the same spatial and temporal frequency filter. The filter’s spatial frequency passband was centered at a spatial frequency of 2.5 cycles per degree and had a bandwidth of 0.5 octaves. Its temporal frequency passband was centered at a speed of 2.5 degrees per second and had a bandwidth of 1 octave. The filter’s orientation bandwidth was 3 degrees. For each stimulus condition, the stimulus set contained five unique filtered noise movies. Each orientation discrimination experiment included stimuli that varied in orientation and contrast. There was one high and one low contrast level per experiment. The high contrast value was constant across experiments, the low contrast value varied somewhat across experiments. The high contrast stimuli spanned a range of 11 different orientations, the number of low contrast orientations varied across experiments (15 experiments had 11 orientations, 7 had 9, 3 had 7, and 4 had 2).

### Fixation task

At the beginning of each recording session, monkeys first performed a passive fixation task. We used a hand-mapping procedure to estimate the location of the spatial receptive fields of visually responsive units. The average receptive field center estimate served as the center location for the visual stimuli presented during the rest of the recording session. We conducted an initial fixation task during which we presented sinusoidal gratings of varying orientation for 1000 ms each. This was followed by the orientation discrimination task.

### Orientation discrimination task

The orientation-discrimination task is a variant of classical visual categorization tasks in which the subject uses a saccadic eye movement as operant response^42,55,56^. We used a richer version of this task in which subjects are invited to additionally report their confidence in each perceptual decision. Each trial began when the subject fixated a small white dot at the center of the screen. Upon fixation, four black choice targets appeared — one in each quadrant of the screen. Targets to the left of the fixation point represented counter-clockwise decisions, targets to the right clockwise decisions. Upper choice targets indicated high decision confidence, lower choice targets low confidence. After a variable pre-stimulus fixation period the stimulus appeared in the near periphery (average eccentricity: monkey F = 4.32^°^, monkey Z = 3.00^°^) for 500 ms. Subjects judged the orientation of the stimulus relative to vertical. The stimulus then disappeared along with the fixation mark and subjects reported their decision and confidence with a saccadic eye movement to one of the four choice targets. Auditory feedback was given to indicate the accuracy of the decision and the chosen level of confidence. Specifically, the tone differed for correct and incorrect trials and the sound was played twice in quick succession for high confidence reports. If the decision was correct, a liquid reward was delivered via a solenoid-operated reward system (New Era). Vertically oriented stimuli received random feedback. Trials in which the monkey did not saccade to one of the choice targets within 3 seconds were aborted. To incentivize meaningful confidence reports, there were four possible reward levels. It required one correct decision to move from level 1 to 2, 3 further correct decisions to move from level 2 to 3, and 3 more to reach level 4. Subjects remained at level 4 until they reported an incorrect decision with high confidence, which reset the score to level 1. The higher the reward level, the larger the reward for a correct decision. In addition, correct decisions reported with high confidence were rewarded more generously than correct decisions reported with low confidence. High confidence rewards for each level were 0.04, 0.16, 0.32, 0.64 ml for monkey F and 0.116, 0.232, 0.464, 0.928 ml for monkey Z. Low confidence rewards were a scalar function of low confidence reward. This scalar value varied across sessions and was adjusted to titrate the proportion of high and low confidence responses (average 0.68 ± 0.04 for monkey Z and average 0.82 ± 0.04 for monkey F. Lower scalar values encouraged more high confidence responses due to a larger reward difference between high and low confidence. Each trial, the current reward level was indicated to the monkey by the duration of the pre-stimulus fixation period (the lower the reward level, the longer this duration). Both monkeys managed to stay at the highest reward level for the majority of trials (fraction of trials at reward level 4: monkey F = 64 %, monkey Z = 80 %). We conducted 12 successful recordings from monkey F and 17 from monkey Z (average number of reward level 4 trials per session, monkey F = 753; monkey Z = 1026).

### Human psychophysical experiment

Nineteen human subjects (10 male, 9 female; ages 19-32) with normal or corrected-to-normal vision participated in the experiment. The human behavioral task was the same as the animals’ orientation discrimination task, with the exception that the stimulus was presented more centrally and subjects earned points instead of liquid reward (points per high confidence correct response as a function of reward level: 2, 4, 8, 16; points per low confidence correct response: 1, 2, 4, 8). Human subjects began by completing 175 training trials. We used these initial trials to estimate each subject’s orientation sensitivity. This sensitivity estimate determined the range of stimulus orientations used in the main experiment. We chose the range such that the subjects’ overall task performance level would resemble that of the animals. This procedure worked well for all but two subjects for whom we discarded the first block of trials. Subjects performed the main task in sub-blocks of 50 trials. Subjects were rewarded with monetary points in the same manner as the macaques were rewarded with liquid reward, and received analogous auditory feedback at the end of each trial. Every 50 trials, subjects were given additional visual feedback on their total point count. Subjects completed three blocks of 1100 trials. Eleven subjects judged the same filtered noise stimuli as the monkeys did, eight subjects were presented with deterministic sinusoidal gratings instead. Because meta-uncertainty did not systematically differ across both groups of subjects, we included all these datasets in our analysis except for the two subjects who had poorly calibrated first blocks yielding a total of 17 human observers in this analysis.

### Behavioral analysis

We measured observers’ behavioral capability to discriminate stimulus orientation by fitting the relationship between stimulus orientation and probability of a ‘clockwise’ choice with a psychometric function consisting of a lapse rate and a cumulative Gaussian function. To compare the behavioral capability associated with low and high confidence reports, each psychometric function had its own steepness parameter (the standard deviation of the cumulative Gaussian). The parameters controlling lapse rate and the point of subjective equality (the mean of the cumulative Gaussian) were shared across both psychometric functions. Model parameters were optimized by maximizing the likelihood over observed data, assuming responses arise from a Bernoulli process. For the analysis documented in Fig. 1c, each dataset was analyzed independently.

For each dataset, we obtained an estimate of the subject’s level of ‘meta-uncertainty’ by fitting the CASANDRE model to the choice-confidence data using a fitting procedure described previously^24^. In brief, the model had eight parameters: the standard deviation of the decision variable (*σ*_*d*_) (one per contrast level, two in total), the decision criterion (*C*_*d*_) (one per contrast level, two in total), the level of meta-uncertainty (*σ*_*m*_), the confidence criterion (*C*_*c*_) (we allowed for choice-dependent asymmetries, two in total), and lapse rate (*λ*). For each dataset, we computed the log-likelihood of a given set of model parameters across all choice-confidence reports and used an iterative procedure to identify the most likely set of parameter values (specifically, the interior point algorithm used by the Matlab function ‘fmincon’). For the analysis documented in Fig. 1d, the first and last block of trials completed by the human subjects were analyzed independently.

### Electrophysiological recordings

During the orientation discrimination task, we recorded extracellular spiking activity from populations of V1 neurons through a chronically implanted recording chamber. Every recording session, we used a microdrive (Thomas recording) to mechanically advance one or two linear electrode arrays (Plexon S- and V-probes; 32 or 24 contacts) into the brain. We positioned the linear arrays so that they roughly spanned the cortical sheet and removed them after each recording session. Continuous neural data were acquired and saved to disk from each channel (sampling rate 30 kHz, Plexon Omniplex System). To extract responses of individual units, we performed offline spike sorting. We first automatically spike-sorted the data with Kilosort^57^, followed by manual merging and splitting as needed (with the ‘phy’ user interface, https://github.com/kwikteam/phy). Given that the electrodes’ position could not be optimized for all contact sites, most of our units probably consist of multineuron clusters. We used the fixation task to identify visually responsive units whose activity selectively depended on stimulus orientation. We measured each unit’s response by expressing spike times relative to stimulus onset and counting spikes within a 1,000-ms window following response onset. For each unit, we chose a response latency by maximizing the stimulus-associated response variance^58^. We visually inspected orientation tuning curves and excluded untuned units from further analysis.

### Linear decoders

To assess how well the recorded populations could support the perceptual task, we trained linear *stimulus decoders* to discriminate between clockwise and counterclockwise stimuli. We used all stimuli whose orientation differed from 0 degrees. We first Z-scored each unit’s spike counts. We then used these z-scored responses to estimate the set of linear weights, **w** = (*w*_1_, …*w*_*n*_) that best separate clockwise and counter-clockwise stimulus response patterns, assuming a multivariate Gaussian response distribution:

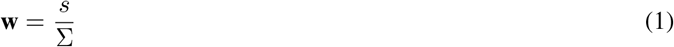

Where *s* is the mean difference of the stimulus-category conditioned Z-scored responses and Σ is the covariance matrix of the Z-scored responses. The decoder weights are calculated from observed trials. To avoid double-dipping, we excluded the trial under consideration from the calculation and solely used all other trials to estimate the weights. This way, we obtained a ‘cross-validated’ stimulus judgement from the linear stimulus decoder for each trial. We quantified how well these decoders captured the animals’ behavior by computing the fraction of consistent perceptual choices and subtracting the number expected by chance based on the decoder’s and the animal’s overall success rate (Fig. 2b). In a later analysis, we compared nonlinear *choice* and *confidence decoders* with their linear counterparts (Fig. 2e,f). For this analysis, we used exactly the same set of training and hold-out trials for the linear decoders as we used for the nonlinear decoders.

### Nonlinear decoders

We trained feed-forward multi-layer perceptron neural networks on Z-scored V1 responses to either predict the animals’ perceptual choice or their confidence report. We implemented networks within the TensorFlow framework using the AdamW optimiser with an objective to minimize binary cross-entropy. Models consisted of 1 hidden layer with 15 hidden units per layer, had a dropout rate between layers of 0.1, and the learning rate was set to 0.001. We explored various hyper-parameter settings and found the results presented here to be robust across settings. We trained networks on 80% of trials (training/validation set) and obtained a cross-validated prediction on the held out 20% of trials, rotating trials between training and held out set such that each trial had a cross-validated prediction (Fig. 2e,f). To ensure that every trial would be part of the hold out set, we trained 30 different networks per dataset. Just like we did for the linear decoders, we solely used cross-validated choice and confidence predictions in our analysis. For each trial, we selected the decoder’s prediction from the “first” network in which this trial was held out. We found 30 networks to be sufficient for every trial to be held out at least once.

We ensured that choice decoders could not use decision confidence to predict choices and that confidence decoders could not use perceptual choice to predict confidence. Specifically, we orthogonalized choice and confidence information in the training trials by maintaining a fixed ratio of high and low confidence reports across clockwise and counterclockwise choices and a fixed ratio of clockwise and counterclockwise choices across high and low confidence reports. To do so, we randomly selected trials from under-represented trial types (e.g “high-confident counter-clockwise”) and concatenated them to the training set. To minimize potentially confounding influences of cross-trial variation in the animals’ motivation, attention, and alertness, we only included “reward level 4” trials in the training set. Training sets on average contained 1088 trials.

To interrogate whether the confidence decoders extracted meaningful information from V1 responses, we compared the slope of the psychometric function conditioned on the confidence decoder’s output (Fig. 3a). This comparison is most reliable when both psychometric functions contain a similar number of trials. We achieved this by using the median of the latent confidence variable as confidence criterion. We did this for both high and low contrast trials.

We probed the confidence decoders with synthetic patterns of neural activity (Fig. 3c). To create these patterns, we first computed the cross-trial average firing rate per unit for a given stimulus orientation using only high contrast, reward level 4 trials. We manipulated the gain of these responses by multiplying this average population response with a scalar factor. For each recording session, we randomly picked on the 30 trained networks for this analysis (Fig. 3d).

We studied the decoders’ latent variables (Fig. 4). This analysis involved computing a discriminability index. To do so, we first Z-scored the latent variables per stimulus condition, thus removing stimulus-driven effects. We then created two groups of trials based on the animals’ behavioral reports (either their perceptual choice or their confidence report). We included all stimulus conditions for which both response options had been used at least 5 times. Finally, we computed the area under the curve for both sets of trials^36^.

## Supplementary Information

**Supplementary Figure 1.**
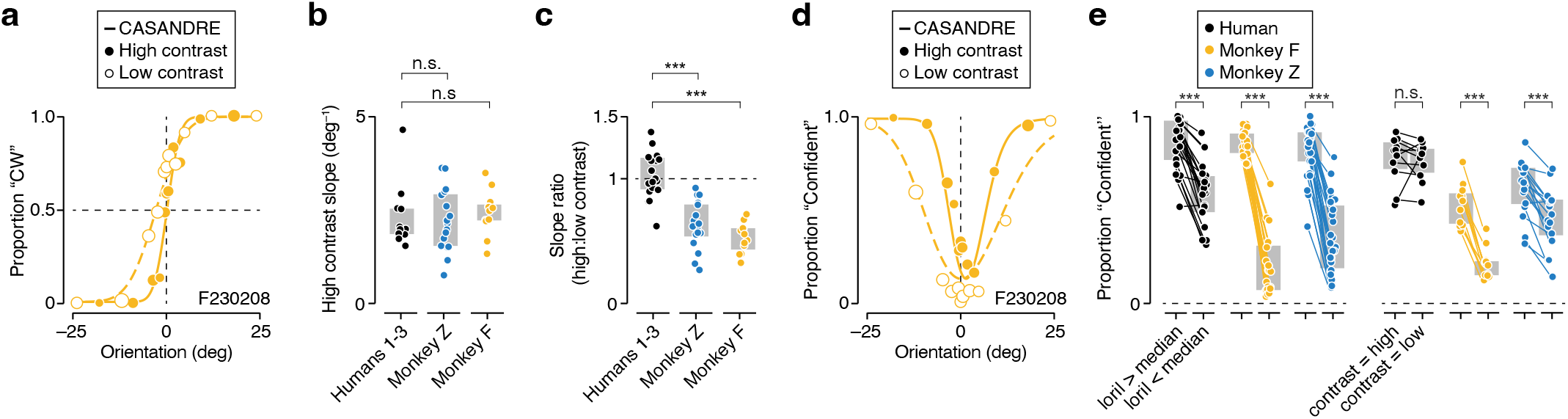
Further comparison of human and monkey behaviour. (**a**) Psychophysical performance for monkey F in an example recording session. Proportion of clockwise (CW) choices is shown as a function of stimulus orientation for high and low contrast stimuli (filled vs open symbols). Symbol size reflects the number of trials (total 1390 trials, slope ratio = 0.46). The curves are fits of a process model for confidence (CASANDRE^24^). (**b**) Analysis of perceptual sensitivity. The slope of the high contrast psychometric function is shown for a group of human observers and both monkeys. We only included the human subjects for whom stimulus contrast and stochasticity was identical to the values used in the monkey experiments (n = 11). Orientation sensitivity of the humans and monkeys was comparable. Grey bars indicate interquartile range. (**c**) The ratio of the slope of the high and low contrast psychometric functions for a group of human subjects and both monkeys. The contrast manipulation had a stronger impact on the monkeys’ perceptual sensitivity. We speculate that this may be due to the stimuli being presented more centrally in the human experiments. Grey bars indicate interquartile range. (**d**) Proportion of high confidence judgments as a function of stimulus orientation for high and low contrast stimuli (filled vs open symbols) in an example recording session. (**e**) (Left) Proportion of high confidence reports elicited by stimuli whose orientation is more or less extreme than the median stimulus orientation. (Right) Proportion of high confidence reports elicited by high and low contrast stimuli. Grey bars indicate interquartile range. n.s. not significant, * *P* < 0.05, ** *P* < 0.01, *** P < 0.001.

**Supplementary Figure 2.**
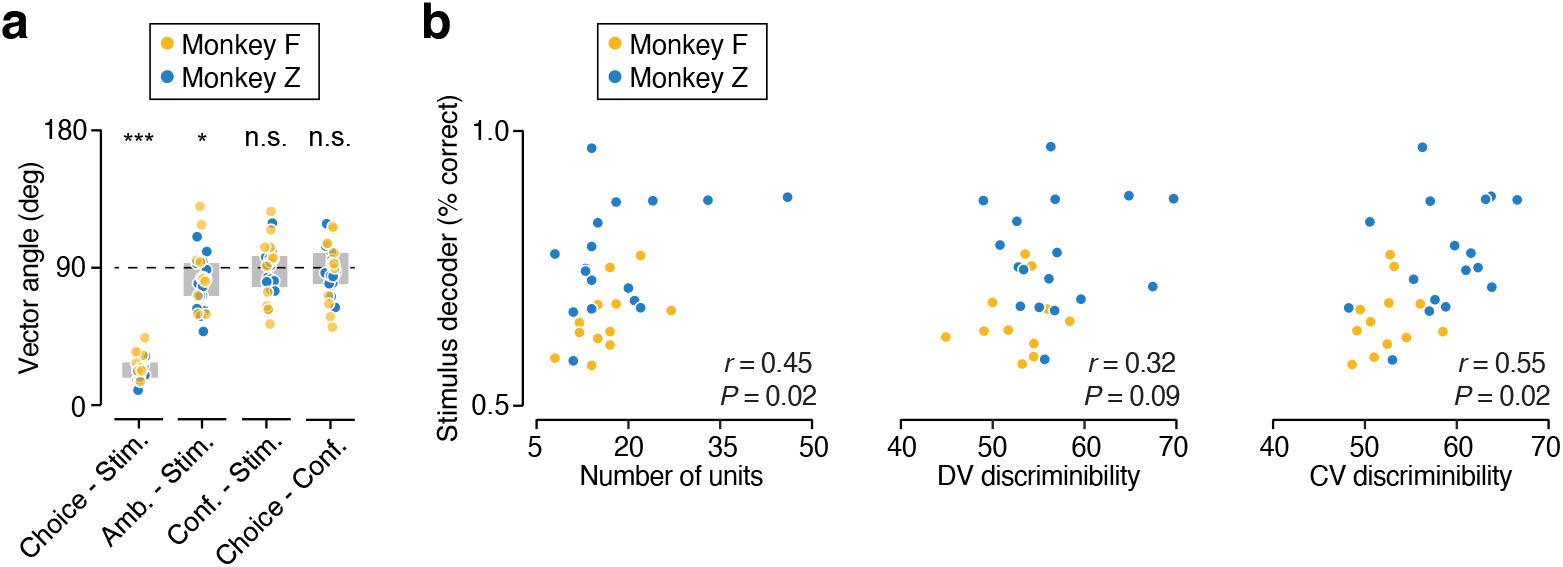
Further analysis of linear and non-linear decoders. (**a**) Comparison of the orientation of the hyper-planes used to slice neural population space by a linear *choice* decoder (trained on all data), *stimulus* decoder (trained on all non-ambiguous trials), *ambiguous* decoder (trained to predict choices on ambiguous trials only), and *confidence* decoder (trained on all data). Unrelated hyperplanes tend to be orthogonal in high-dimensional spaces (vector angle = 90 deg). Note that this is not the case for the choice and stimulus decoder, nor for the stimulus and ambiguous decoder. Grey bars indicate interquartile range. (**b**) Task performance of the linear stimulus decoder is plotted against the number of units in the population (left), the non-linear choice decoder’s latent variable’s discriminability (middle), and the non-linear confidence decoder’s latent variable’s discriminability (right). Larger neural populations tended to be better able to support the perceptual task. Populations that contained more stimulus information tended to enable better decoding of perceptual choices and confidence reports. n.s. not significant, * *P* < 0.05, ** *P* < 0.01, *** P < 0.001.

